# Dynamic adjustment of feeding behavior depending on dominant-subordinate relationship in large-billed crows

**DOI:** 10.64898/2025.12.21.694083

**Authors:** Kazutaka Morita, Ei-Ichi Izawa, Hiroshi Matsui

## Abstract

Animals that forage socially adjust their behavior according to social status. Dominance and subordination are defined by priority of access to resources in competitive contexts. While dominant individuals can control access to resources, subordinates must obtain food through alternative strategies. In direct encounters with dominants, subordinates are expected to adjust their behavior to the partner on a moment to moment basis. However, the decision dynamics in such situations remain unclear. We tested large-billed crows in a dyadic feeding experiment to examine whether social status shapes feeding behavior. Using tracking analysis, we quantified the time series of approach velocity toward food and assessed whether its dynamics were modulated by the partner’s movements. Subordinates adjusted the velocity of their reaching movements. Specifically, a dominant’s approach increased a subordinate’s reaching speed, suggesting that subordinates accelerated to avoid aggression under dominance pressure. In contrast, this effect was weak or absent in dominants, indicating asymmetric social influences on foraging. Taken together, these findings suggest that subordinates fine tune the kinematics of feeding behavior in response to dominant movements, whereas dominants do not.

## Introduction

In social species, individuals adjust their behavior in accordance with dominance and subordination. Considerable attention has been paid to the formation and function of dominant–subordinate relationships (Cheney & Seyfarth, 2007; Drews, 1993; Hasegawa & Kutsukake, 2015; Hasegawa et al., 2015; Miyazawa et al., 2020; Sapolsky, 2005). Effects of conspecific presence have been documented across domains and taxa (e.g., chicks: Ogura & Matsushima, 2011; humans: Markus, 1978; macaques: Santos et al., 2012; rats: Sekiguchi & Hata, 2019; fishes: Dzieweczynski et al., 2005; Wong et al., 2008; roosters: Shimmura et al., 2015; calves: Vázquez-Diosdado et al., 2025).

In social groups, dominance and subordination are defined by consistent outcomes for a dyad across successive contests (Drews, 1989) and are often linked to competition over food (Abbott & Dill, 1989; Brouns & Edwards, 1994; Ekman & Lilliendahl, 1993; Tamm, 1977). For example, Tamm (1977) found that high-ranked jackdaws (*Corvus monedula*) tended to drive lower-ranked individuals away from feeding sites, indicating that dominant individuals gain energetic benefits on average. By contrast, some studies report that subordinates obtain food under pressure from dominant conspecifics by employing non-aggressive alternatives such as stealing (Adams et al., 1998; Gomez-Melara et al., 2021; Schmidt & Hoi, 1999). Indeed, quantitative analyses show that subordinates can still secure sufficient amounts of food (Hollis et al., 2004; Kidjo et al., 2016). However, it remains unclear which aspects of behavior animals adjust to obtain food as a function of their position in the dominant and subordinate relationship. Because dominance is relative rather than absolute and varies with the specific opponent, animals may need to alter their behavior flexibly in response to momentary social circumstances. For instance, dominant individuals may be less likely to display aggression toward subordinates while engaged in feeding. These considerations motivate experimental investigations that quantify momentary behavioral adjustments as a function of the particular conspecific encountered.

Corvids are well suited to address this question because members of the family exhibit notable behavioral flexibility across social, motor, and cognitive domains (Clayton & Emery, 2015; Güntürkün & Bugnyar, 2016). Several corvid species form fission–fusion aggregations and range widely during prolonged juvenile periods of three to four years, including ravens (*Corvus corax*), carrion crows (*C. corone*), and large-billed crows (*C. macrorhynchos*) (Boucherie, Loretto, Massen, & Bugnyar, 2019;

Coombs, 1978; Izawa, 2011; Loretto et al., 2017; Uhl et al., 2019). In large-billed crows, the present subjects, stable dominance–subordination relationships are consistently observed (Izawa & Watanabe, 2008; Miyazawa et al., 2020). In addition, their feeding behavior is guided by momentary visual information (Matsui & Izawa, 2017, Matsui & Izawa, 2019ab). Large-billed crows are therefore ideal subjects for investigating feeding behavior in social contexts.

To quantify how dominance–subordination influences momentary feeding behavior, it is necessary to measure dynamic features of behavior over time, because any behavior consists of a series of movements. Recent advances in movement tracking enable scrutiny of subtle, moment-to-moment modulations of behavior (Anderson & Perona, 2014; Mathis & Mathis, 2020). Therefore, in addition to the traditional measure of the number of food items obtained, we quantified kinematic changes in feeding behavior, specifically the velocity profile, as an index of the effects of a conspecific. Specifically, in our setting this denotes instances in which the dominant and the subordinate approach the food. Pecking in crows typically occurs within one second (Matsui & Izawa, 2019b). Because partner effects are difficult to record manually in real time, an approach that combines image-based tracking with statistical modeling is advantageous.

Taken together, this study aimed to test whether crows adjust their behavior during feeding according to immediate social circumstances defined by dominance–subordination relationship. Specifically, we compared the movement velocity of feeding behavior between dominant and subordinate individuals to assess whether the influence of a conspecific varies with social status. To this end, we developed a food scrambling experiment in which a dominant and a subordinate were positioned face-to-face across a food arena. Feeding behavior was recorded with high-speed camcorders to capture momentary movements, and the influence of the partner’s behavior on movement was quantified using time-series modeling. Three predictions were considered. First, if a conspecific exerts an effect irrespective of social status, the effects should not differ between dominants and subordinates. Second, the effect should be absent if crows do not attend to the behavior of the other individual. Third, if subordinates adjust their behavior in the presence of dominants, the influence of the conspecific should be asymmetric, such that subordinates are more affected by dominants, whereas dominants are not similarly affected.

## Material and methods

### Subjects

Twelve male juvenile or adult large-billed crows (*Corvus macrorhynchos*) were used. The experiment was conducted in 2018. All birds were captured in accordance with local government regulations (Permit no. 4005). Sex was genetically determined from blood using polymerase chain reaction. Crows were individually housed in stainless-steel mesh cages (W 43 cm × L 60 cm × H 50 cm), separated by 15 cm to prevent the establishment of dominance relationships through direct interactions outside the experiment. They could see and hear one another but had no physical contact during experimental periods. Birds were assigned to three groups of four based on year of capture (before 2014, 2015, or 2016). Within each group, all six possible dyads were tested, yielding a total of 18 dyads. Food was withheld for three hours before daily experimental trials to establish motivation, and sufficient food in cups was provided after each trial. Water was available ad libitum in the home cages. The room was maintained at 21 ± 2 °C on a 13L:11D cycle with lights on at 08:00. Subjects were given regular access to a large outdoor aviary (W 150 cm × L 280 cm × H 170 cm) to reduce stress from spatial constraints. All housing and experimental procedures complied with Japanese National Regulations for Animal Welfare and were approved by the Animal Care and Use Committee of Keio University (no. 14077 and 20119).

### Determination of dominance relationships

To determine dominance relationships within each dyad, we used the dyadic encounter procedure from our previous study (Izawa & Watanabe, 2008). Dominance between the two individuals in each dyad was verified by five successive pairwise encounters. Each individual participated in one encounter per day.

In each encounter, two individuals were tested to determine their relative dominance in a food competition context. At the start of the encounter, the experimenters gently released the two birds into an experimental cubicle (W 150 cm × L 170 cm × H 120 cm) with an interval of approximately 20 s. One minute after the encounter began, a transparent plastic bag containing seven pieces of dog food was dropped onto the center of the floor to promote social interaction. The encounter ended when all food was consumed or when five minutes had elapsed. Each encounter was video recorded from above the cubicle for offline behavioral analyses.

Winners and losers in each encounter were identified from social behaviors. The loser was the only individual that exhibited submissive behavior, such as begging vocalizations, fluffed head feathers, or avoidance (Izawa and Watanabe, 2008). The other individual was coded as the winner. Encounters were scored as draws when both individuals showed submissive behavior or when neither did. Dominance within a dyad was assigned from the outcomes of five encounters. An individual that won all five encounters was designated the dominant, and its opponent was designated the subordinate. A dyad was classified as tied when neither bird achieved more than two wins. Only dyads with a clearly established dominant and subordinate were included in subsequent experiments, because the purpose was to test the effects of a stable dominance relationship on movement during social foraging.

### Food scrambling task

#### Apparatus

Two chambers (W 45 cm × L 45 cm × H 120 cm) faced each other across a food stage (W 45 cm × L 25 cm) in an experimental room (Figure 1). The side of each chamber facing the food stage was a transparent front panel. Each panel had an 8 cm gap through which crows could peck food on the stage while preventing them from stepping onto it. To allow simultaneous access, small gates were installed in front of both gaps. The gates were controlled by an Arduino microcontroller, enabling the experimenter to open them from outside the experimental room. The food stage contained 21 holes (0.7 cm diameter) arranged in seven rows and three columns. Cheese was cut into columns and placed in the holes. Using this apparatus, we recorded both crows’ movements synchronously from a horizontal view during the experiment.

**Figure 1.**
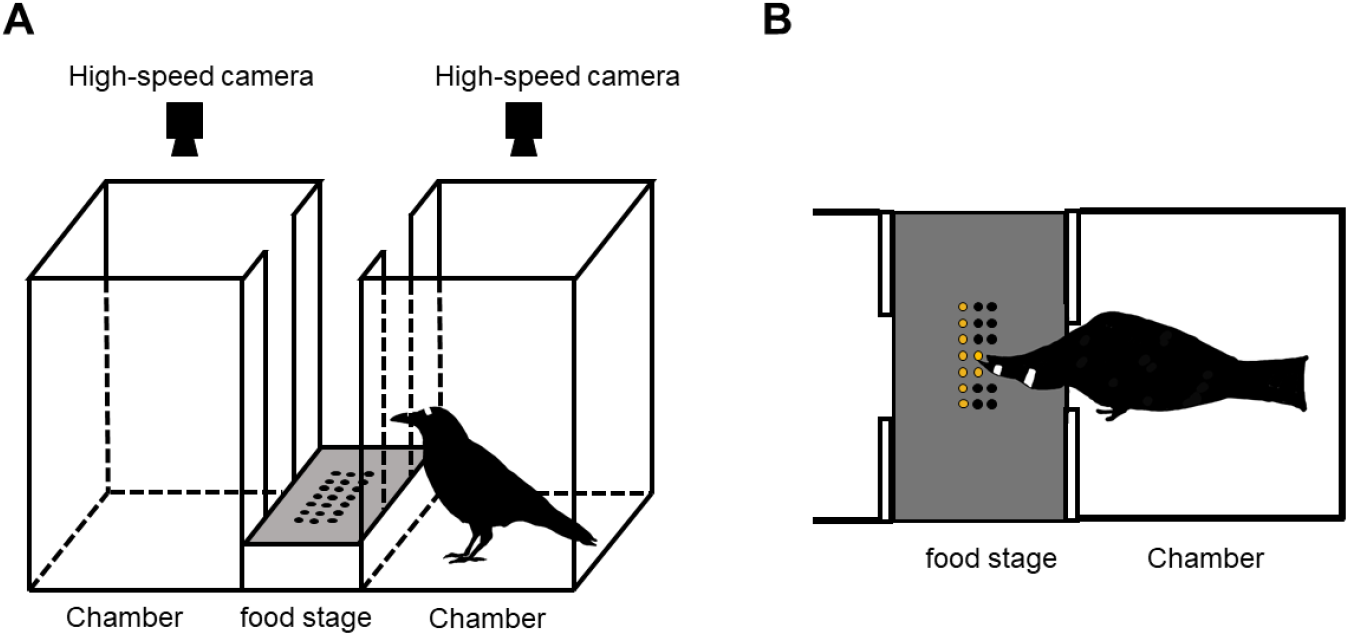
(A) Schematic illustration of the apparatus used in the food scrambling task. Two chambers faced each other across a food stage. The side of each chamber facing the food stage was fitted with atransparent stop panel containing a narrow gap through which the crows could peck at food items placed on the stage, while preventing them from stepping onto it. Two cameras were positioned above the chambers, allowing synchronized video recordings of both crows’ movements from a horizontal view during each trial. The gate located in front of the gap on the stop panel is omitted from this illustration. (B) Schematic illustration of the horizontal view of the apparatus and a crow consuming food. The food stage contained 21 holes. The gate in front of the gap on the stop panel is omitted from this illustration.

#### Procedures

The experiment comprised four phases: habituation and training, a pre-control (single-crow) session (five trials per subject), a scrambling session (five trials per dyad), and a post-control (single-crow) session (five trials per subject). In the habituation phase, a crow was released into each chamber for 10 minutes on three consecutive days, with no food available on the food stage. The gates to the food stage remained closed to prevent direct interaction. In the subsequent training phase, one crow was placed in the experimental chamber, and seven pieces of cheese were made available in one column of holes on the food stage. Training continued until the crow consumed all seven pieces. This procedure was conducted three times for each of the three columns to confirm that crows could consume cheese from all positions on the stage.

After training, each crow underwent the pre-control session, in which a single crow was tested to verify motivation for food. The pre-control session was followed by the scrambling session, in which a dyad was placed on opposite sides of the apparatus to quantify the effect of dominance on feeding movements. Finally, the post-control session, identical to the pre-control session, was conducted. Crows performed one trial per day, and trials were run either daily or on alternate days. Each trial consisted of presenting 21 pieces of cheese arranged on the food stage and began when the gates were opened, allowing access to the stage. Trials ended after three minutes. The side of the chamber assigned to each crow was quasi-randomized.

#### Video recording and tracking of crow’s movements

Movements were recorded with two high-speed cameras (120 frames per second; CASIO EXILIM EX-ZR70, Japan) mounted above each chamber of the experimental apparatus (Figure 1). A brief LED flash was delivered immediately after recording began to enable synchronization of the videos during post hoc analysis. Removable white tracking markers (rectangular adhesive tape) were attached to the upper beak and transversely across the head. From the horizontal view, we extracted two-dimensional x–y coordinates frame by frame using the Python-based software DeepLabCut (Nath, Mathis, Chen et al., 2019). The tracked points were the center of the beak marker and the left and right edges of the head marker, and these coordinates were used to compute the kinematic variables described below.

### Analysis

#### The number of obtained food in the scrambling task

Because the dominance–subordination relationship had been established in advance, dominants were expected to have priority access to food. In our task, however, 21 pieces of cheese were presented to the birds, allowing subordinates occasional opportunities to snatch food when dominants were distracted. To examine whether the number of food items obtained differed between dominants and subordinates, we used a generalized linear mixed-effects model (binomial distribution, logit link). Dominance status (dominant or subordinate) was included as a fixed effect, and individual identity was treated as a random effect. The significance of independent variables was assessed using a likelihood ratio test at the 5% significance level.

#### Vector autoregressive model

Time-series analyses were conducted for each feeding bout, defined as the period beginning when an individual initiated movement to obtain food and ending when food acquisition was completed. Each feeding bout was manually identified from the videos by the experimenter. The extracted coordinates were smoothed with a smoothing spline to reduce noise prior to analysis. To characterize dynamic interactions between dominant and subordinate individuals, we applied a bivariate vector autoregression (VAR) model (Figure 2). The model was originally introduced in macroeconometrics to describe mutually dependent time series (Sims, 1980), and have been applied to neuroscience (Zhang, Liu, Ji, & Huang, 2017), human child–mother interaction (Chen, Chow, Hammal, Messinger, & Cohn, 2021; Hoerzer, Liebe, Schloegl, Logothetis, & Rainer, 2010), and animal behavior (Ewing, Riggs, & Ewing, 2007; Wang et al., 2025). We briefly describe the model here, and more formal description appears in Supplementary Material.

**Figure 2.**
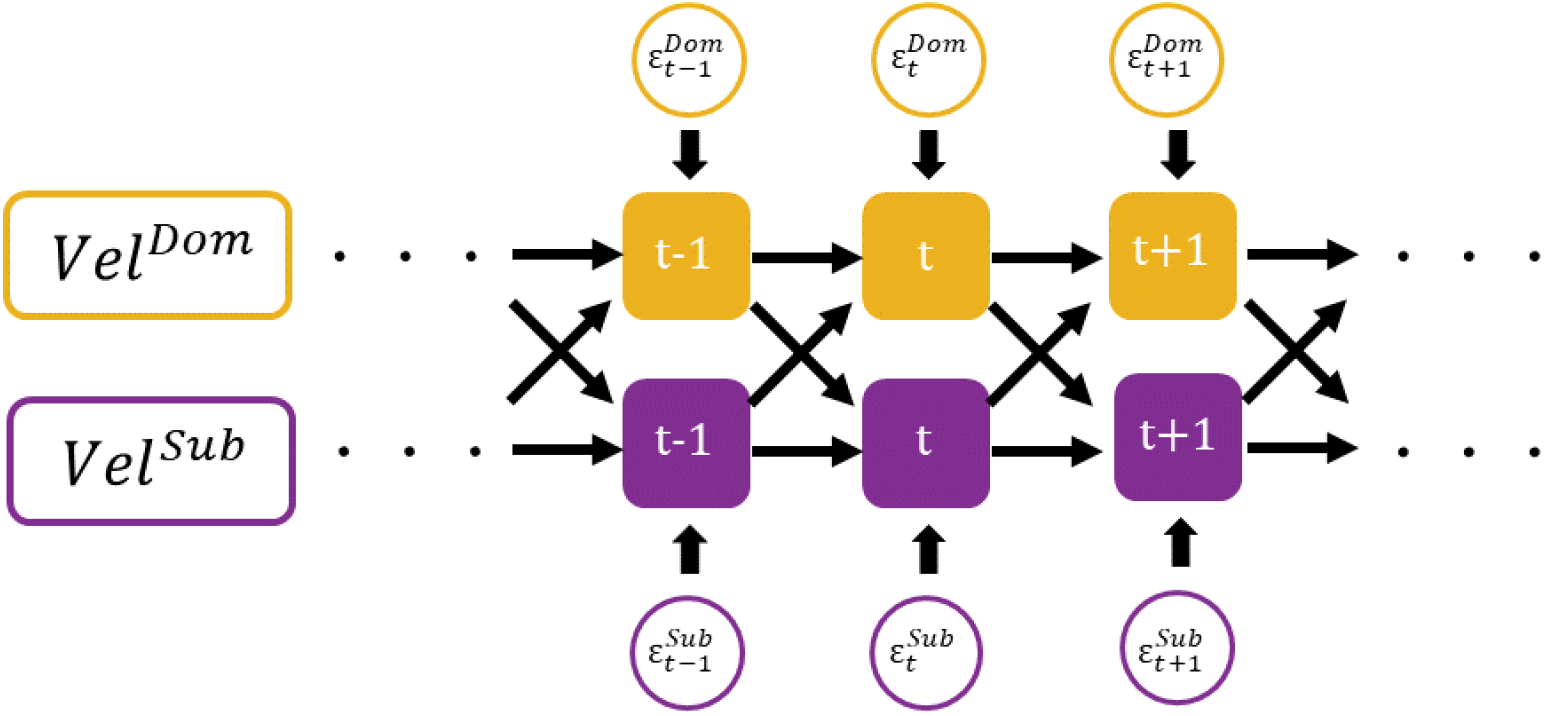
Schematic diagram of the VAR (Vector Autoregressive) model used in the analysis. . The time series of velocity for the dominant and subordinate individuals in each trial are denoted as *Vel*^*Dom*^ and *Vel* ^*sub*^, respectively. *t* represents the time step, set at 8 ms according to the temporal resolution of the camera. Black arrows indicate the coefficients between time steps. In the VAR model, each variable is influenced by its own lagged values, the past values of the other variable, and an error term. Accordingly, in our model, *Vel* ^*Dom*^ at time t is influenced by *Vel* ^*Dom*^ and f*Vel* ^*sub*^ at time t-1, as well as the error term 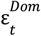 . Similarly, *Vel* ^*sub*^ at time t is influenced by *Vel* ^*sub*^ and *Vel*^*Dom*^ at time t-1, and error term 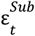 . This model thus quantifies the mutual influence of movement between the dominant and subordinate individuals.

The VAR model assumes that two or more time series evolve based on three factors: their own history, their mutual interaction, and random error terms (Figure 2). In our case, a dyad of crows produces head movements during feeding behavior, represented as velocity (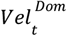 and 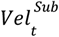). Naturally, temporal variation in this velocity is influenced by the individual’s own recent past, since an animal cannot instantly accelerate from zero to its maximum speed. In the model, this dependence is captured by regression coefficients linking past data points of the same individual to its current value (the autoregressive model; horizontal arrows in Figure 2). In addition, an individual’s movement may be affected by the motion of its dyadic partner. Such interaction effects are estimated as regression coefficients from the partner’s recent data points to the individual’s current velocity (diagonal cross arrows in Figure 2). Through these time-series regressions, the VAR model evaluates the dyadic influence between individuals.

Given the large number of parameters estimated in a VAR model, interpreting individual coefficients is difficult. Therefore, after fitting the VAR model to both crows’ movements, we computed impulse response functions (IRFs). An IRF simulates how a shock to one variable affects another using the fitted model. This approach is widely used to examine the dynamics of VAR models (Ewing et al., 2007; Sims; 1980; Swanson & Granger, 1997). Conceptually, an IRF represents the derivative of the forecast path with respect to a one-unit increase in the error term. We denote the impulse response of subordinates to changes in dominants as *IRF*^*sub*→*Dom*^, and that of dominants to subordinates as *IRF* ^*Dom*→*sub*^. In short, an IRF can be interpreted as the effect of one individual’s movement on the other’s movement. We applied the VAR model and calculated IRFs for each trial of dyads during the scrambling session. This yielded a pair of IRFs per trial— *IRF*^*Dom*→*sub*^ and *IRF*^*sub*→*Dom*^—resulting in 74 IRF pairs in total (five trials from 15 dyads, with one trial excluded due to mechanical error).

#### Bayesian modeling for time series analysis

Using the IRF analysis, we obtained *IRF*^*Dom*→*sub*^ and *IRF*^*sub*→*Dom*^ for each trial. Our main question was whether the temporal profiles of these IRFs exhibited an asymmetry in influence between dominants and subordinates, and if so, when such a difference emerged. To test this, we employed a state-space modeling framework, which is widely used in time-series analysis (see Commandeur & Koopman, 2007, for a tutorial). In brief, we introduced a difference parameter *diff*_*t*_ between the two IRFs and modeled how *diff*_*t*_ varied over time (see Supplementary Material for details). By performing Bayesian estimation of both the intrinsic temporal changes and the dynamics of this difference, we evaluated whether a significant divergence existed between the two IRFs. Statistical significance of difference of state space-model in the parameters *diff*_*t*_ can be inferred when 95% credibility intervals (CI) of posterior distribution do not overlap zero.

The state-space model was implemented via the probabilistic language Stan (Carpenter et al., 2016). Markov Chain Monte Carlo methods (MCMC) were used for parameter estimation. All iterations were set to 10,000, burn-in samples were set to 2000, and thinning samples were set to 4, with the number of chains set to four. The convergences of the models were assessed by checking Gelman-Rubin convergence statistic (R-hat, Gelman and Rubin, 1992) of all parameters under 1.1, and visually inspecting the trace-plot.

All analysis was performed with R 3.5 including the packages ‘tidyverse’ (Wickam, 2017), ‘vars’ (Pfaff, 2008), and ‘tseries’ (Trapletti, &Hornik, 2019).

## Results

Dominant–subordinate relationships, defined by social behavior (aggressive and submissive displays), were successfully established in all 15 dyads, consistent with previous reports in large-billed crows (Izawa & Watanabe, 2008; Miyazawa et al., 2020). In both the pre-control and post-control phases (single-individual feeding), dominant and subordinate individuals each consumed most of the available food (pre-control: mean = 87%, SE = ±0.03; post-control: mean = 75%, SE = ±0.06).

In the scrambling phase, we fit a generalized linear mixed model with food consumption rate as the dependent variable. The amount of food consumed differed significantly between dominant and subordinate individuals, consistent with the expected function of dominance–subordination (likelihood ratio test, *X*^*2*^ = 0.30, p < 0.01; Figure.3). Nevertheless, unlike prior observations in which dominants frequently monopolized food, subordinates also consumed a substantial portion in our food-scrambling task, obtaining 43% ± 2% SE on average. Therefore, subordinates also succeeded in obtaining a substantial amount of food by taking advantage of moments when the dominants were inattentive.

**Figure 3.**
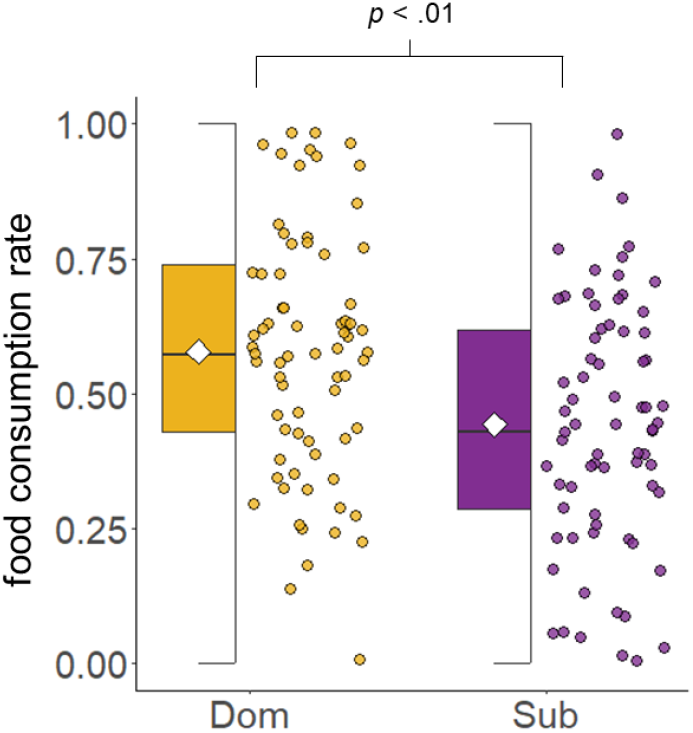
Food consumption rates of dominant and subordinate crows in each trial during the scrambling phase. A value of 1 indicates that the crow obtained all 21 pieces of cheese placed on the food stage of the apparatus. The white rhombus in each box plot represents the mean value for each group.

To quantify the mutual influence of movements associated with the partner’s behavior, we applied the VAR model and calculated the IRFs of both individuals’ velocities in each trial (Figure 4). The hand, *IRF* ^*sub*→*Dom*^ indicates the movement of dominant crow approach to food stage will cause increased velocity of subordinate crow’s approach to food, and the effect was at maximum 0.53 s. On the other *IRF* indicates the less profound effect, which was at maximum 0.33 s. The curves of IRF obtained from VAR model seems that asymmetric effects between dominants and subordinates on their movements of feeding behavior. To verify the difference between *IRF* ^*Dom*→*sub*^and *IRF* ^*sub*→*Dom*^ over a time, we applied state-space model described above, and estimated the time-varying of parameter*diff*_*t*_, resulting in the significantly higher *IRF*^*Dom*→*sub*^ than *IRF*^*sub*→*Dom*^ from 0.33 s (the gray bar in Figure 4). The impulse response began to diverge approximately 0.3 seconds after the partner’s movement, suggesting that subordinates rapidly detected the dominant’s behavior and adjusted their own movements accordingly.

**Figure 4.**
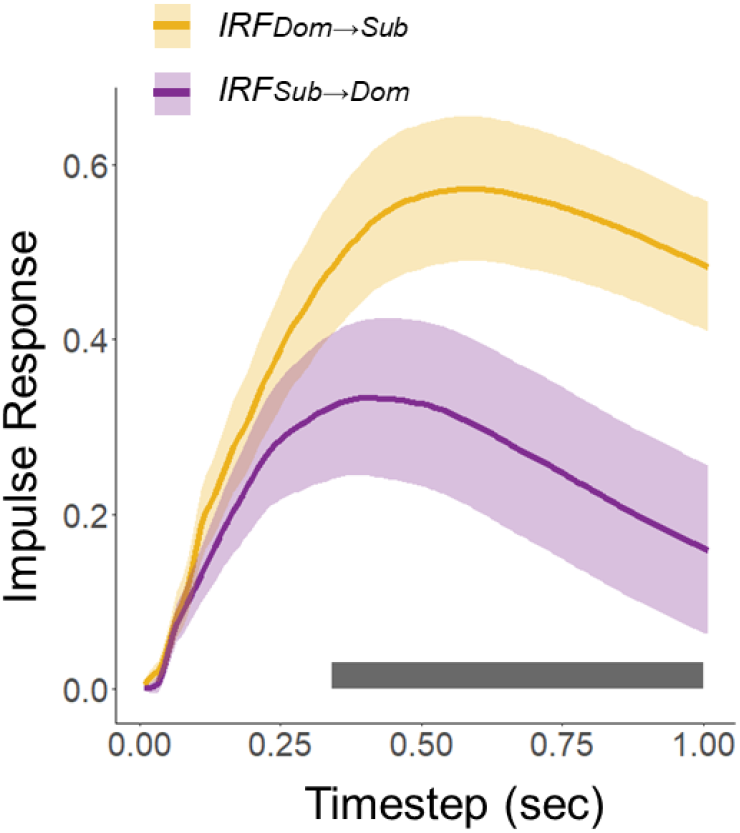
Difference between *IRF*^*Dom*→*sub*^and*IRF*^*sub*→*sub*^. The solid line represents the mean, and the shaded area represents the standard error for each group. Positive velocity values indicate an increase in velocity toward the food stage (approach), while negative values indicate a decrease in velocity away from the food stage (withdrawal). The gray bar with an asterisk marks time steps showing a significant difference, where the 95% credible interval of the posterior distribution for the time-varying parameter diff[t] in the state-space model does not overlap with zero.

## Discussion

This study tested whether crows adjust their momentary movements in response to those of a conspecific. To this end, we developed a food scrambling task in which two crows faced each other across a food arena. Consistent with prior work in crows (Izawa and Watanabe, 2008), dominants obtained more food than subordinates. However, in our task, dominants did not monopolize food, and subordinates obtained a moderate amount despite the risk of aggression. These results are not unexpected; rather, they are useful for elucidating the behavioral strategies by which subordinates pilfer food from dominants.

Time-series analysis revealed asymmetric effects, with subordinates strongly influenced by the movements of dominants, whereas dominants showed little sensitivity to subordinate movements. Because dominants can displace subordinates by attacking when a subordinate approaches food, they may not need to attend closely to subordinate behavior. By contrast, subordinates must employ nonaggressive strategies to obtain food under dominance pressure. Consequently, subordinate crows did obtain food in our experiment, although in smaller amounts than dominants.

In social animals, the frequency of aggressive behavior often increases with higher social rank. (Adams et al., 1998; Miyazawa et al., 2020), and so aggression likely represents a primary behavioral repertoire of dominants. By contrast, nonaggressive tactics for obtaining food under competition have been reported in more naturalistic settings (Adams et al., 1998; Manson & Appleby, 1990; Schmidt & Hoi, 1999; Taillon & Côté, 2007). For example, “sneaking” or “stealing” denotes a subordinate tactic in which an individual snatches food when aggression can be avoided (Adams et al., 1998; Hollis et al., 2004; Schmidt & Hoi, 1999). Field observations confirm that subordinates can secure moderate amounts of food without attacking dominants (Hollis et al., 2004; Kidjo et al., 2016).

In our results, the influence of dominants on subordinates was expressed with a temporal delay of approximately 0.5 s. This delay may reflect subordinate tactics that reduce the risk of aggression from dominants. During feeding, crows approach the food, make contact, and then withdraw the head to swallow; attacking may be difficult during swallowing. In our data, when time zero was defined as first contact with the food, swallowing began approximately 0.3 to 0.4 s thereafter. Thus, subordinates withheld movement and initiated an approach after the dominant began to move toward the food. Comparable moment-to-moment adjustments have been reported in monkeys. Subordinate monkeys generally refrain from taking pellets located near dominants, but when dominants temporarily cease to display signs of dominance, subordinates reduce submissive behavior and begin to take food (Chadwick-Jones, 1998; Delgado, Delgado-García Amérigo, & Grau, 1975; Santos et al., 2012). Although dominance establishes priority of access to resources, subordinates dynamically adjust their behavior to momentary social circumstances. In this study, specifically, subordinate crows modulated their feeding kinematics as a function of the dominant’s movement velocity.

Such behavioral adjustment to a conspecific’s actions may be linked to social characteristics such as hierarchical structure and the degree of fission and fusion dynamics (Aureli et al., 2008; Amici et al., 2013, Amici et al., 2018; Boucherie, Loretto, Massen, & Bugnyar, 2019). By definition, social rank is a relative metric, and most individuals, apart from those at the very top or bottom of the hierarchy, must alter their behavior depending on the opponent they face. Individuals of intermediate rank, in particular, regularly encounter both higher- and lower-ranked partners. Moreover, in societies with fission and fusion dynamics where social structures fluctuate over time, individuals may need to infer the dominance status of unfamiliar conspecifics from their observed relationships with familiar ones (Paz-y-Miño et al., 2014). Crows also recognize individuals using visual and auditory cues (Kondo et al., 2012). Together, these observations suggest that crows can utilize the relative status of other individuals to guide their own behavior. Consistent with this view, our results indicate that crows flexibly modulate the velocity of their feeding movements in response to the behavior of conspecifics.

In conclusion, this study tested whether crows adjust feeding behavior according to the social status of the opposing individual. The results showed that, first, dominants obtained more food than subordinates, consistent with previous findings, yet dominants did not fully monopolize access. Subordinates secured a moderate proportion of the food. Second, movement analyses indicated that subordinates sensitively adjusted momentary movement velocity, whereas dominants did not. In particular, subordinates increased approach velocity when dominants moved toward the food. This adjustment likely contributes to feeding success under the risk of aggression from dominants.

## Acknowledgements

This research was supported by JSPS KAKENHI (Grant Number 16J04383 to H.M., 19K21818, 20H01787 to E.I.) and JST CREST (JPMJCR17A4 to E.I.), and Keio University Grant-in-Aid for Innovative Collaborative Research Projects (MKJ1905 to E-I).

## Declaration of generative AI

The authors used ChatGPT 5.0 for language-proofing to refine the manuscript written by non-native English speakers. After using this tool/service, the authors reviewed and edited the content as needed and takes full responsibility for the content of the publication.

